# Water soluble polysaccharides from *Spirulina platensis* exhibited cyto/DNA protection and suppress the growth of gastric cancer cells via modulation of galectin-3

**DOI:** 10.1101/2021.12.16.473001

**Authors:** Vinayak Uppin, Shylaja M Dharmesh, R Sarada

## Abstract

Polysaccharides from natural sources play a significant role in the management of different cancer types including gastric cancer. In this study, we reported the effect of spirulina polysaccharide (Sp) on galectin-3 modulatory activity on gastric cancer cells. The polysaccharide was isolated from the spirulina biomass, characterized, and the *in silico, in vitro* studies are carried out to assess the bioactivities. The isolated Sp possessed average molecular weight of 1457 kDa, and galactose (42%) as major sugar along with Rhamnose, Arabinose, Xylose, and Mannose. Further, characterization of Sp by FT-IR and NMR spectrum indicated the presence of (β1-4D) galactose sugar with galactoarabinorhamnoglycan backbone. Among the monosaccharides, galactose showed highest binding affinity with galectin-3 protein as evidenced by the *in silico* interaction study. The obtained Sp, inhibited the proliferation of AGS gastric cancer cells by 48 % without affecting normal NIH/3T3 cells as opposed to doxorubicin, a known anticancer drug. Also, Sp exhibited galectin-3 mediated haemagglutination inhibition with MIC of 9.37 μg/mL compared to galactose 6.25 μg/mL, sugar specific to galectin-3. The Sp treatment significantly (p<0.05) lowered the expression of galectin-3 by 32 % compared to untreated control cells. In addition, Sp exhibited the potent cytoprotection in RBCs, Buccal cells, and DNA exposed to oxidants. Thus, the findings suggest that the polysaccharide from spirulina offer a promising therapeutic strategy in the management of gastric cancer in addition to its currently known nutritional and pharmaceutical applications.

## 1 Introduction

Metastasis is the most critical and complex step in cancer, which involves cell adhesion, cell invasion, angiogenesis, proliferation, and inhibition of cell death, ultimately leading to dysregulated cell growth and formation of aggregated tumours. Over the past several decades, cancer is continued to be leading cause of mortality across the world [Fitzmaurice, et al., 2017]. However, the complex multistep events involved in metastasis are the result of the interplay of multiple molecules like galectin-3. The human galectin-3 is a member of galectin family consisting of carbohydrate recognition domain (CRD), and N terminal short peptide which is widely expressed on most cellular surface of many tissues [Yang, et al., 2008, Fortuna-Costa et al., 2014]. The role of galectin-3 in metastasis is well depicted in our previous studies [Balasubramanian, et al., 2009], as well as others [Takenaka, et al., 2002; Funasaka, et al., 2014; Song, et al., 2014]. Since gastric cancer is the second leading cause of mortality worldwide [Ferro, et al., 2014], and is closely associated with dietary factors, any aberrations in dietary pattern can lead to an imbalance in pro/antioxidant status, increased metastasis, angiogenesis, and dysregulation of apoptosis. The present study therefore focused on isolation of spirulina polysaccharide (Sp), elucidation of structural, chemical composition, and evaluation of galectin-3 modulatory activity in addition to its cytoprotective properties in the human gastric adenocarcinoma cells (AGS).

Further, the efforts are directed towards identifying a potent naturally occurring molecule with bioactive properties like gastro protective, cytoprotective, and anti-metastatic activity. Among the naturally occurring biomolecules the food-derived molecules and microalgae like spirulina are well documented for health beneficial effects from ancient times [Kay, et al., 1991; Matos, et al., 2017]. Spirulina (*Spirulina platensis*), a blue-green algae, is garnering the attention of several researchers across the globe due to its nutritional composition, bioactive molecules having varied applications in food, feed, nutraceutical, cosmetic and therapeutic industries [Seyidoglu, et al., 2017]. With the advancement in the knowledge of algal science, spirulina is being used as a nutraceutical food supplement due to its high protein and vitamin content. Among the several phytochemicals present in spirulina, the bioactive studies are mainly focused on carotenoids, polyphenols or phycocyanin [Park, et al., 2018; Gabr, et al., 2020]. Meanwhile, the use of dietary polysaccharides in preventing/treating complex diseases is in the log phase [Ahmadi, et al., 2017; Pynam, et al., 2019; Besednova, et al., 2020], and many research groups are exploring the algal sources for the dietary polysaccharides. Although, polysaccharides are produced from many biological sources such as seaweed, plants, fungi, and algae, they differ in structure and composition, which decides their bioactivities. Polysaccharides, particularly from different algal sources such as *Arthospira, Gracilaria*, and *Ulva*, gained so much attention because of their special physicochemical properties and high biological activities thus considered as marine drugs [Han, et al., 2017; Patil, et al., 2018]. The previous studies reported are limited to the antioxidant properties of water soluble and sulfated polysaccharides from spirulina [Chaiklahan, et al., 2013] and the claims on anticancer properties were not explored in cancer cells [Kurd, et al., 2015].

The current study therefore, focused on the following aspects (1) Isolation of water-soluble polysaccharides from spirulina biomass, (2) Characterization for the sugar composition, molecular weight, and structural backbone, (3) Cyto/DNA protection of Sp upon exposure to oxidants in RBCs, buccal cells, and DNA, (4) *In silico* interaction of monosaccharides of Sp with galectin-3, and (5) Whether inhibition of galectin-3 result in inhibition of gastric cancer cells growth; If so, what will be its effect on the normal cells.

## 2 Materials and Methods

### 2.1 Chemicals

Ethidium bromide, butylated hydroxyanisole (BHT), gallic acid, calf thymus DNA, acridine orange, ruthenium red and HPLC grade carbohydrate standards such as rhamnose, arabinose, xylose, mannose, galactose, and glucose, protease, etc., were purchased from Sigma Chemical Co, St. Louis MO, USA. The folin-ciocalteu’s phenol reagent, and hydrogen peroxide, from Merck Ltd (Mumbai, India). Ham’s F-12, MTT (3-[4,5-dimethyl-2-thiazolyl]-2,5-diphenyl 2-H-tetrazolium bromide) (TC-191), were procured from Hi-media, India (Cell Culture grade). The Anti-galectin-3 monoclonal (ab2785) antibodies were procured from Abcam, UK. The other chemicals such as Agarose, trypsin, EDTA (ethylene diamine tetra acetic acid), Kojic acid, calf thymus DNA, hexane, sodium phosphate buffer, glutaraldehyde, sodium chloride, Tween twenty, TEMED, SDS, amberlite IR 120H+ resin, sulphuric acid, and solvents used were of the analytical grade purchased from Sisco Research Laboratories, Mumbai, India.

#### 2.2 Isolation and Physicochemical characterization of Sp

*Isolation of Sp from spray dried powder:* The spray dried spirulina biomass powder was obtained from the department of plant cell biotechnology, and the polysaccharide isolation was carried out with slight modification over the protocols of Phatak, et al., 1988 [Phatak, et al., 1988]. Briefly, the powder was defatted by soxhlet apparatus with hexane in the ratio of 200 ml/g (v/w). The defatted sample (100 g) was subjected to specific enzymatic digestions (protease, termamylase and glucoamylase) to remove proteins, amylase and amylopectin by providing optimum reaction conditions for enzymes. Then, the deproteinized sample was extracted using soxhlet apparatus with water without using solvent. The solution was filtered and the supernatant was subjected to precipitation with four volumes (v/v) of absolute ethanol at 4°C. After 3 hours the pellet was separated by centrifugation at 5000 g at room temperature for 20 min to at least three times. Further, the pellet was washed twice with 50 mL of absolute ethanol, resuspended in 10 mL of water to get the water soluble fraction of polysaccharide, and lyophilized to obtain water soluble spirulina polysaccharide (Sp). The yield of the polysaccharide was calculated by the equation

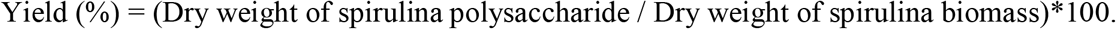

##### Determination of purity and molecular weight

The purity of Sp was analyzed by cellulose acetate electrophoresis (*CAE)* as per the method described by Wegrowski, & Maquart, 2001 [Wegrowski, et al., 2001]. The polysaccharide sample was dissolved in deionized water and loaded onto the membrane at 10 μg/mL and 20 μg/mL. Then the membrane is stained with ruthenium red, the image is documented and observed for the migration of molecules, sharpness and intensity of the band. The average molecular weight (Mv) of Sp was quantified by the intrinsic viscosity method. Briefly, Sp solution [1 % in 0.5 % acetic acid solution] was used for the viscosity plot generation, and Mv was calculated by Mark-Houwink equation (η) = K_m_Ma where, k_m_ =3.5×10^−4^ and a=0.76 as constants for the acetic acid solution [Rinaudo, et al., 1993].

##### Sugar composition analysis by gas-liquid chromatography (GLC)

The monosaccharides of Sp was quantified by GLC according to the method of Sathisha, et al., 2007 [Sathisha, et al., 2007]. Briefly, 10 % sulphuric acid was added to 10 mg of Sp, and subjected to acid hydrolysis for 8-12 h. Later, the sample was treated with sodium barium carbonate and amberlite IR 120H^+^ resin, for neutralization and deionization respectively. Then, the Alditol acetates were prepared and subjected to sugar composition analysis with RTX-1 column using a Shimadzu GLC (Kyoto, Japan). The flow rate was kept at 40 ml/min, the column temperature adjusted to 200 °C, and the injector temperature was maintained at 250 °C.

##### FTIR and NMR analysis

Sp was subjected to IR spectral study using a Perkin-Elmer spectrum 2000 spectrometer (Connecticut, USA) and the absorbance mode was at a resolution of 4 cm^-1^ with a wave number range 4000-400 cm^-1^. The Sp sample of 10 mg was dissolved in 1000 μL deuterated water and subjected to ^1^H and ^13^C NMR analysis. The NMR spectrum was recorded using the Bruker AQS 400 MHz NMR spectrophotometer. The ^1^H NMR spectrum was recorded at 500 MHz of 10,330 Hz spectral width involving water presaturation pulse program zgpr. Similarly, the ^13^C spectrum was recorded at 125 MHz with a spectral width of 28,985 Hz.

##### Total sugar, uronic acid, total phenols, and calcium content in Sp

Total sugar content of Sp was estimated by the phenol-sulphuric acid method [Rao, et al., 1989]. To various aliquot of extracts, 300 µL of 5 % phenol and 2 mL of concentrated H_2_S0_4_ was added. Absorbance was measured against a blank at 490 nm with glucose as a reference standard. Likewise, the uronic acid estimation was carried out as described previously [Bitter, 1962]. Briefly, to a different aliquot of Sp, 3 mL of concentrated H_2_S0_4_ was added. The mixture was boiled for 20 min and temperature was brought to 25 °C and carbazol (0.1 %) was added. Then, the mixture was incubated for 2 h in the dark and absorbance was measured at 530 nm. The concentration of uronic acid in samples was estimated against galacturonic acid as a standard. Further, the total polyphenol content in the Sp powder was determined by Folin Ciocalteu’s (FC) phenol reagent method [Swain, et al., 1959]. The aliquots of Sp, with standard gallic acid, were employed in the assay. The total volume was made to 1mL using distilled water; to this, 2 mL of Na_2_CO_3_ (10 %) and 1mL of FC reagent (1:1 v/v) were added. The reaction mixture was incubated at ambient room temperature in the dark for 1 h, and absorbance was measured at 765 nm. The standard graph was generated using gallic acid (5 to 25 μg/mL), and the concentration of phenol content in the Sp was calculated. The calcium content in the Sp is estimated by using the methods and protocols supplied by the kits [Agappe kit – 11006001].

### 2.3 Cyto and DNA protective assays

#### RBC’s protection assay

The RBCs were isolated according to the procedure described by Ashadevi et al., 2014 [Ashadevi et al., 2014]. In brief, 20 μL of RBCs were incubated with Sp aliquots in 20mM phosphate buffered saline (PBS), pH 7.4, in the presence of oxidants (H_2_O_2_), and the total volume was adjusted up to 1000 μL with buffer. The reaction mixture was incubated at 37 °C for 30 min and centrifuged at 1500 *g* for 10 min. The haemolysis was spectrophotometrically evaluated at absorbance of 410 nm, and the cell pellets were fixed in 3% glutaraldehyde on a cover slip and dehydrated using increasing concentration of acetone (30–100 %). Then the samples were mounted onto an aluminum stub (100–200 A^0^) and the cells were observed under a scanning electron microscope (SEM) and images were taken at magnification (×10000).

#### Buccal cells protection assay

The cellular damage was induced on normal buccal cells (NBCs) by exposing cells to UV radiation at different time intervals and validated by dye staining such as acridine orange and ethidium bromide. The modulatory effect of Sp was compared to gallic acid upon exposure of cells to UV followed by the treatments. NBCs were scraped from the buccal cavity and added to the PBS (pH 7.4), centrifuged at 1500 *g* for 10 min. This step was repeated until the supernatant exhibit no absorbance at 280 nm, and the 10 μL Sp (5 mg/mL) was added to the 50 μL PBS suspension of cells. Then, the reaction mixture was incubated at 37°C for 10 min, followed by exposure to UV at 10, 20, and 30 min time intervals. The Acridine orange + ethidium bromide dye (10 μL) was added to visualize the changes in cells and the images were photographed at a magnification of 40X and documented. The live cells stained green with acridine orange and red with ethidium bromide dyes [Baskic, et al., 2009].

#### DNA protection assay

The *in vitro* DNA protective effect of Sp was electrophoretically determined using calf thymus DNA [Von Gadow, et al., 1997]. The reaction mixture (20 μL) contained a calf thymus DNA, PBS [20 mM/L, pH-7.4], and various concentrations of Sp samples, was incubated for 10 min at 37 °C. The hydrogen peroxide (oxidant) was then added, incubated for 2 h at 37°C and mixed with bromophenol blue dye and electrophoresis was performed on 1 % agarose gel. The relative migration difference between the native, oxidized, Sp treated DNA was observed on 1 % agarose gel after staining with ethidium bromide (Et-Br), and gel documentation (Syngene G: Box, Haryana, India).

### 2.4 *In vitro* anticancer activity study

#### Galectin-3 inhibition assay

The Sp was evaluated for its galectin-3 inhibitory activity based on hemagglutination assay. This assay was carried out with the slight modifications over the protocols described by Gao, et al., 2013. Briefly, agglutination assay was carried out in a microtiter V plate and the urine sample from the cancer patients was used as a source of galectin-Each well of the plate contained a reaction mixture of 25 µL of 1% BSA, 25 µL of Sp or 0.15 M NaCl (negative control), 25 µL of 1.5 µg/mL Gal-3 (urine), and 25 µL of 4% (v/v) human erythrocyte suspension. The PBS buffer (0.1 M, pH 7.0) is used to make the dilutions and incubated at 4 °C for 90 min for the agglutination. The minimum inhibitory concentration (MIC) of the Sp, and the standard galactose was determined by the serial dilutions.

#### Cell culture

The AGS human gastric carcinoma cells and NIH-3T3 normal mouse embryonic fibroblast cells were obtained from the National Centre for Cell Sciences (NCCS) Pune, India. The cells were cultivated in Ham’s F-12 and DMEM medium respectively containing L-glutamine, sodium bicarbonate, and supplemented with 10 % fetal bovine serum (FBS) at 37 °C. T-25 flasks are used to grow the cells, regulated with 5 % CO_2_ incubator, 95 % humidity (Eppendorf), and when the cells were confluent washed with PBS (pH 7.4) several times and harvested upon treatment with 0.25 % trypsin. Further, the AGS cells were used for an anti-proliferative study.

#### Galectin-3 expression

The expression of galectin-3 and inhibitory effect of Sp was further determined by western immunoblotting. AGS cells were seeded into 60 mm petridishes (2×10^5^cells/dish) overnight treated with Sp 40 μg/mL, (dose was fixed after MTT assay) and incubated for 48 h. The cells were lysed with RIFA buffer after harvesting with trypsin and the protein content was quantified using the protocol of Lowry’s et al., 1951. Then, the equal quantity of protein (40 µg) was resolved on 10 % polyacrylamide gel and transferred onto the PVDF membrane supplied by Bio-Rad, USA. Next, the membranes were blocked with 5 % skim milk and probed for galectin-3 (1:1000, Abcam, Cambridge, UK) and GAPDH (Cloud-Clone Corp., TX, USA) overnight at 4 ^0^C. Membranes then incubated with appropriate secondary antibodies with intermittent washing. After completing the washing steps, the membranes were treated with enhanced chemiluminescent reagents obtained from Bio-Rad and documented for densitometry.

#### Antiproliferative assay

Antiproliferative activity of Sp, on AGS cells was carried out by MTT assay with slight modifications over the protocols described by Sathisha, et al., 2007 [Sathisha, et al., 2007]. After 12 h of cell growth (2×10^4^ cells/ well), media was replaced with media containing varying concentrations of Sp (10, 20, and 40 μg/mL) and a drug doxorubicin (as a positive control) and cells were incubated for another 48 h. Following incubation, 15 μL of 5 mg/mL MTT solution [5-3-(4, 5-dimethylthiazol-2-yl)-2, 5-diphenyltetrazolium bromide] freshly prepared was added and media was removed after 4 h of incubation, without disturbing the formazan crystals. The formazan crystals formed as a result of reduced product of MTT was solubilised by adding DMSO (100 μL) and absorbance was taken by the microtiter plate reader (Infinite pro, USA), after 20 min, at 570 nm. The results were expressed as percent cell viability for the control, doxorubicin and Sp treatments.

#### The effect of Sp on the induction of apoptosis

The ability to induce apoptosis upon Sp treatment was evaluated using ethidium bromide (Et-Br) and acridine orange dye as described by Jayaram et al., 2015 [Jayaram, et al., 2015]. The stained images of AGS cells were photographed by inverted fluorescence microscope for the qualitative analysis of dead to live cells. Further, the apoptotic activity of Sp on AGS cells was qualitatively evaluated by DAPI (4,6-diaminidino-2-phenylindole) staining. Briefly, the AGS cells were cultured on dishes overnight (2×10^5^ cells/dish) and treated with Sp (20 mg/mL) and drug doxorubicin, incubated for 48h. After harvesting, the cells were washed with phosphate buffer saline (PBS) and stained with DAPI for 30 min at 37°C in the dark and observed under microscope with UV excitation wavelength of 300-500 nm. The cells that exhibited the nuclei with condensed chromatin or fragmented nuclei were considered as apoptotic [Jayaram, et al., 2015].

#### Tyrosinase inhibitory activity

Tyrosinase inhibitory activity of Sp was studied using L-Dopa as a substrate as per the protocols of Rao, et al., 2013 [Rao, et al., 2013]. The inhibitory concentration (IC_50_) of Sp required to inhibit the tyrosinase activity was quantified using standard tyrosinase inhibitor - kojic acid.

### 2.5 *In silico* docking study

The 3D structural coordinates of monosaccharides such as Galactose [ID, 439353], Arabinose [66308], Rhamnose [25310], Mannose [65127], were retrieved from the PubChem database, and the ligand preparation was carried out using Auto-Dock Tools 4.2. The X-ray diffraction-based crystal structure of target protein galectin-3 was downloaded from RCSB-PDB in complex with an inhibitor, and a resolution of < 3.0 A^0^ was selected for the study (PDB ID: 5E88/6FOF). The protein preparation was done by removing water molecules, adding polar hydrogen’s, adding kollman united atom charges, and gasteiger charges to the ligand. The complexes bound to the protein receptor molecule were removed, and molecular docking was carried out using Auto-Dock 4.2. The lamarckian genetic algorithm (LGA) method was followed to find the optimal conformation of the ligand, grid box dimensions were kept minimal, covering the proteins’ active site residues, and grid maps were generated. Using LGA, molecular docking study was carried out, and ten possible outcomes were generated, and the one with the lowest binding energies picked up from the cluster. The visualization of the docked complex files was performed by UCSF chimera, and a 2D interactive map of the complex was generated using Lipgplot+ software.

### 2.6 Statistical analysis

The normality of distribution and homogeneity of variance were checked before analyzing for one-way ANOVA (non-parametric) and significant differences were set to p<0.05, followed by Tukey’s test using Graph Pad Prism version 7.04. The P-value < 0.05 was considered statistically significant and only the mean and standard deviation are plotted in the figure.

## 3 Results

### 3.1 Isolation and physicochemical characterization of Spirulina polysaccharide

*Polysaccharide yield, Carbohydrate, uronic acid, and total phenolics content in Sp:* The polysaccharide yield obtained by soxhlet hot water extraction followed by further purification indicated an enriched yield of 4 %. The content of carbohydrates, polyphenols, and uronic acid content is given in the Table 1. The results indicated the presence of 5.19 mg GAE/g of total phenolics and 735 mg/g of total carbohydrates, and the uronic acid content of 120 mg/g of total sugar.

**Table 1.**
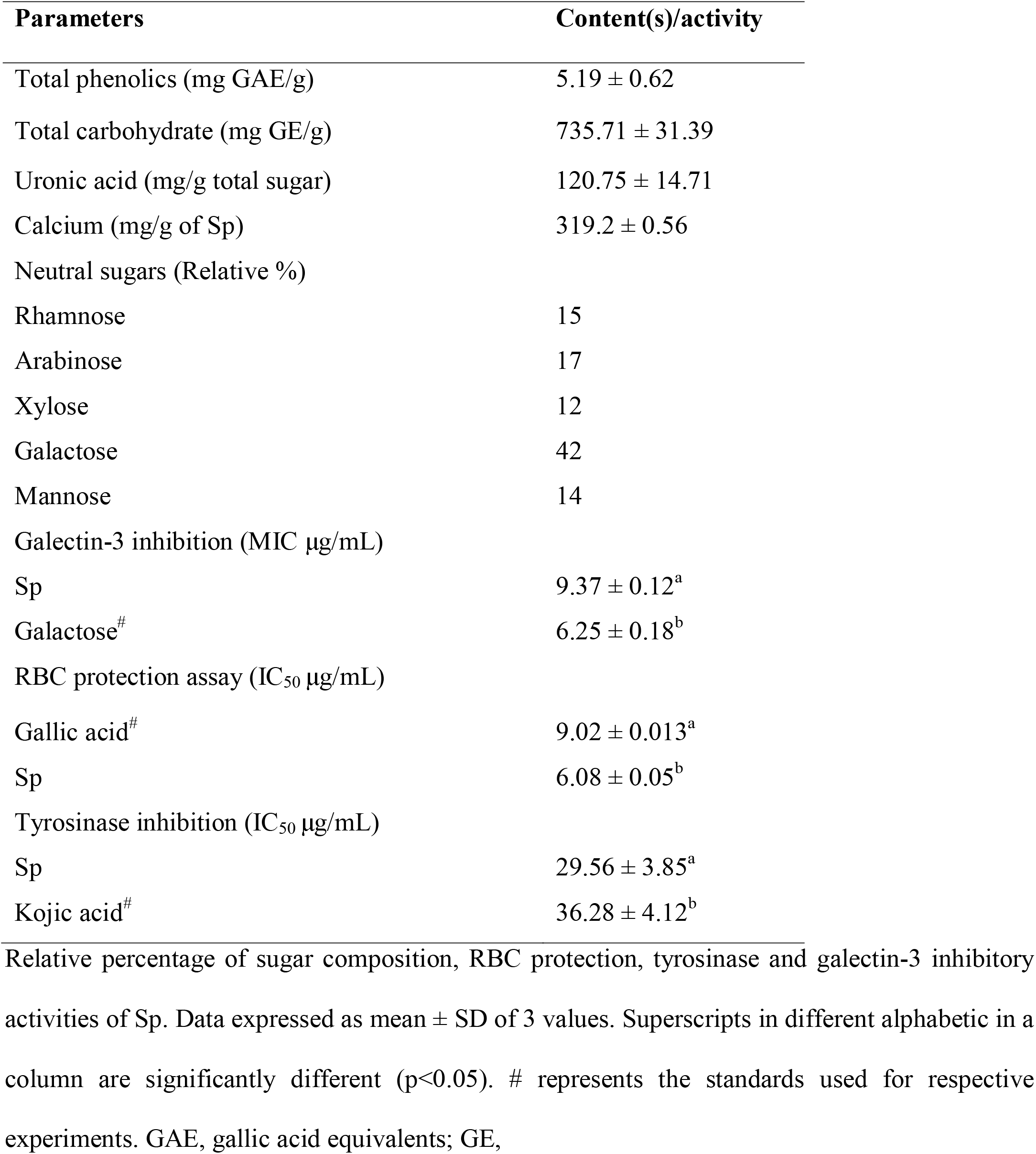
Biochemical composition and bioactivities of Spirulina polysaccharide.

| Parameters | Content(s)/activity |
| --- | --- |
| Total phenolics (mg GAE/g) | 5.19 ± 0.62 |
| Total carbohydrate (mg GE/g) | 735.71 ± 31.39 |
| Uronic acid (mg/g total sugar) | 120.75 ± 14.71 |
| Calcium (mg/g of Sp) | 319.2 ± 0.56 |
| Neutral sugars (Relative %) |  |
| Rhamnose | 15 |
| Arabinose | 17 |
| Xylose | 12 |
| Galactose | 42 |
| Mannose | 14 |
| Galectin-3 inhibition (MIC µg/mL) |  |
| Sp | 9.37 ± 0.12<sup>a</sup> |
| Galactose<sup>#</sup> | 6.25 ± 0.18<sup>b</sup> |
| RBC protection assay (IC<sub>50</sub> µg/mL) |  |
| Gallic acid<sup>#</sup> | 9.02 ± 0.013<sup>a</sup> |
| Sp | 6.08 ± 0.05<sup>b</sup> |
| Tyrosinase inhibition (IC<sub>50</sub> µg/mL) |  |
| Sp | 29.56 ± 3.85<sup>a</sup> |
| Kojic acid<sup>#</sup> | 36.28 ± 4.12<sup>b</sup> |

Relative percentage of sugar composition, RBC protection, tyrosinase and galectin-3 inhibitory activities of Sp. Data expressed as mean ± SD of 3 values. Superscripts in different alphabetic in a column are significantly different (p<0.05). # represents the standards used for respective experiments. GAE, gallic acid equivalents; GE,

#### Purity, Molecular weight, and Sugar composition

The single intense band was observed from the cellulose acetate electrophoresis of the Sp at concentrations (10 µg/mL and 20 µg/mL), (fig. 1B) and this band was scraped and employed for further studies. The average molecular weight as determined by the Mark-Houwink equation revealed that Sp was 1457 kDa, which is in close proximity as reported by Rajasekar, et al., 2019 [Rajasekar, et al., 2019]. The monosaccharide composition analysis, by gas chromatography revealed that Sp is composed of rhamnose (15 %), arabinose (17 %), Xylose (12 %), Mannose (14 %), and galactose (42 %) (Table 1). The monosaccharide galactose was found to be the predominant neutral sugar observed, followed by arabinose and rhamnose. This higher galactose content in the Sp followed by arabinose suggests the presence of galactoarabinournan backbone. Also, the relatively higher amount of rhamnose sugar indicated the possibility of side rhamnournan side chain. The remaining monosaccharides such as xylose and mannose may be originated from either other cell components.

**Figure 1.**
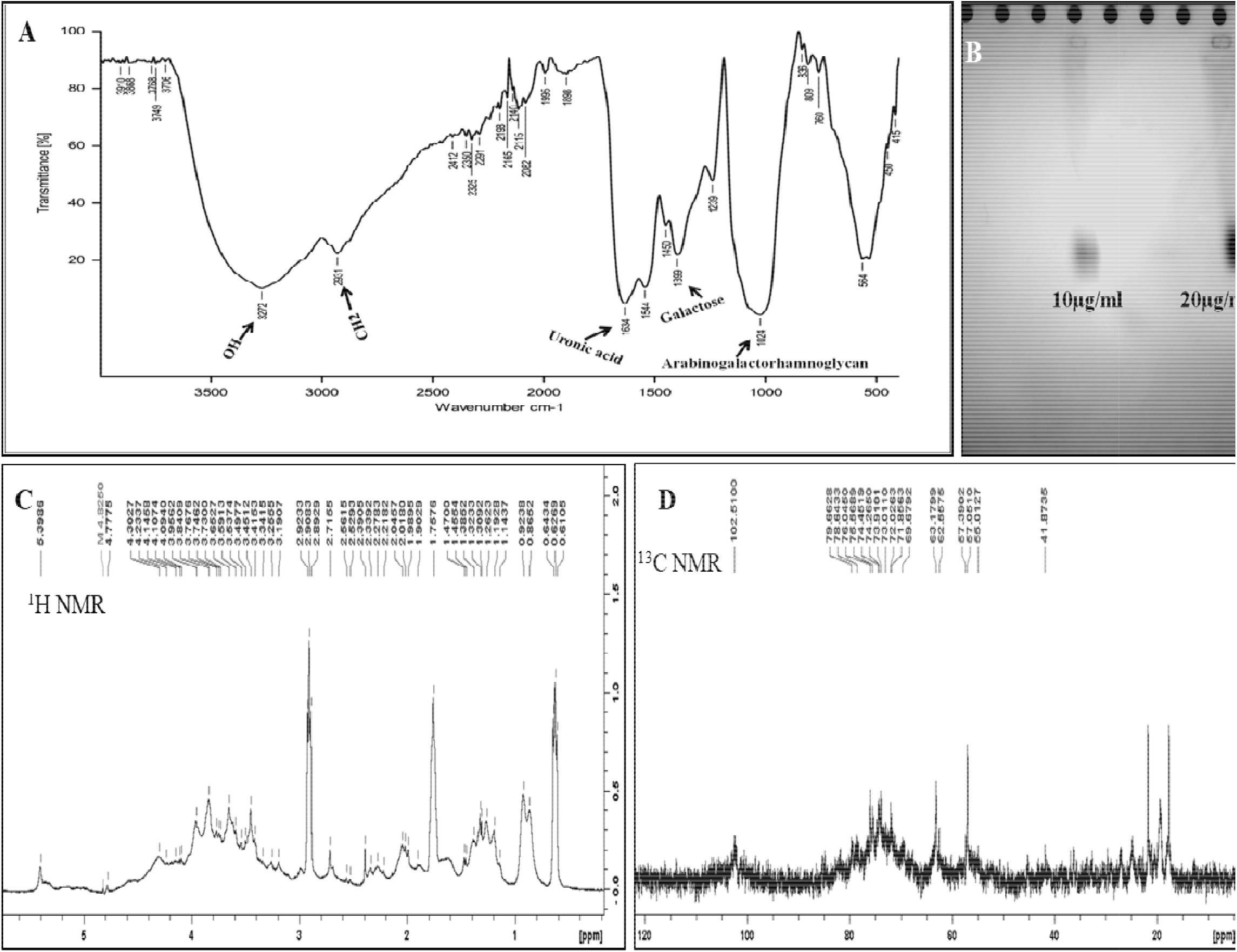
Structural characterization and purity analysis of spirulina polysaccharide (Sp). (A) FTIR spectra of Sp, (B) Cellulose acetate electrophoresis of Sp, (C) ^1^H-NMR and (D) ^13^C-NMR spectra of Sp.

#### Structure elucidation by FT-IR and NMR study

FT-IR analysis of Sp was performed by infrared spectrophotometer, and the spectrum concerning the band position and intensities are presented in Fig 1A and Table 2. The band in the region 3272 cm^-1^ signifies the stretching of the OH group, and 2931cm^-1^ suggests the stretching of CH_2_ group [Peng, et al., 2012]. Likewise, the wave numbers 1634, 1450 cm^-1^ represent the presence of uronic acid content [Boulet, al., 2007], and the wavelength of 1024 cm-^1^ represents the Rhamnogalacturonan or Arabinogalactorhamnoglycan [Harsha, et al., 2016; Kacurakova, et al., 2000]. Further, the IR signal at 1544 cm^-1^ indicated the phenolics content in Sp sample and the wave number 1399 cm^-1^ suggests the presence of relatively high galactose/ β-galactan ratio in the Sp [Coimbra, et al., 2008], which is further supported by neutral sugar composition (Table 1). Similarly, the NMR data revealed that ^1^H NMR peaks in the range between 3 to 5 ppm denote the distinctively presence of polysaccharides. Also, the downstream ^1^H NMR signals between 4.5 to 5.7 ppm indicates the presence of polysaccharides. Further, the chemical shifts of Sp were identified in comparison with the literature reports of dietary polysaccharides (Table 2). The data clearly indicated that our findings are in line with the earlier reports for the NMR chemical shifts of various dietary and algal related polysaccharides [Mallikarjuna, et al., 2018; Lai, et al., 2010].

**Table 2.** FTIR spectra frequencies (cm^-1^) and NMR chemical shifts (ppm) of Spirulina polysaccharides. The FTIR spectral frequencies of Sp are compared with literature.

| FTIR | Sp | Reference |  |  |  |  |
| --- | --- | --- | --- | --- | --- | --- |
| frequencies (cm<sup>-1</sup>) |  |  |  |  |  |  |
| 3272 | Stretching of OH group | Peng et al. (2012) |  |  |  |  |
| 2931 | Stretching of CH<sub>2</sub> group | Peng et al. (2012) |  |  |  |  |
| 1634, 1450 | Uronic acid | Boulet et al. (2007) |  |  |  |  |
| 1544 | Phenolics | Taboada et al. (2010) |  |  |  |  |
| 1399 | Galactose/ β-galactan | Coimbra et al. (1999) |  |  |  |  |
| 1239 | Galactouronic acid (GalpA) | Cui et al. (2007) |  |  |  |  |
| 1024 | Rhamnogalacturonan/<br>Arabinogalactorhamnoglycan | Harsha et al. (2016), Kacurakova<br>et al. (2000) |  |  |  |  |
| 836 | β-D Xylose | Peng et al. (2012) |  |  |  |  |
| <sup>1</sup>H-NMR and <sup>13</sup>C-NMR chemical shifts (ppm) of Sp |  |  |  |  |  |  |
| Residue | C<sub>1</sub>/H<sub>1</sub> | C<sub>2</sub>/H<sub>2</sub> | C<sub>3</sub>/H<sub>3</sub> | C<sub>4</sub>/H<sub>4</sub> | C<sub>5</sub>/H<sub>5</sub> | C<sub>6</sub>/H<sub>6</sub> |
| α-Galactouronosyl | 102/5.7 | 69.6/3.65 | 71.8/3.59 | 71.4/- | 73.9/4.7 | 57.5/3.74 |
| -4)-α-D-GalA-(1- (a) |  |  |  |  |  |  |
| α-Rhamnosyl | - | 76.04/4.1 | 63.1/2.5 | 72.08/2.9 | 73.01/1.7 | -/1.14 |
| 2)- α-L-Rhap-(1 (b) |  |  |  |  |  |  |
| β -Galactosyl | - | 76.5/3.95 | 69.6/3.34 | 71.85/3.41 | 78.64/3.65 | 57.3/3.73 |
| 4)-β-D-Gal-(1 (C) |  |  |  |  |  |  |
| α-Arabinosyl | - | 79.6/3.49 | 73.01/- | -/3.41 | 57./3.2 | - |
| -5)α-L-Ara(1- (d) |  |  |  |  |  |  |

### 3.2. Molecular interaction between Sp monosaccharides and galectin-3

Since Sp had enriched amount of β1-4D galactose, which is a specific sugar that binds to galectin-3, blocking this interaction of galectin-3 mediated metastasis plays a vital role [Balasubramanian, et al., 2009]. Thus, we focused to examine whether Sp binds to galectin-3 protein by virtue of its high β1-4 galactose content. The molecular docking between the monosaccharides and the target protein galectin-3 reveals an appropriate ligand (monosaccharides) that fits both energetically and geometrically to the protein’s binding site. Among the monosaccharides tested for the binding affinity to galectin-3, galactose showed the highest binding affinity with the lowest binding energy (−3.08 Kcal/mol) and dissociation constant (5.50 mM). The binding affinity, dissociation constant (Ki), and amino acid residues involved in the hydrogen/hydrophobic interactions are given in table 3. The Fig. 2 depicts the binding pocket site and the interaction map of Sp monosaccharides with the target protein galectin-3. This data clearly indicated that galactose is the major sugar interacting with galectin-3 with high binding affinity among the list. Although, galactose showed highest binding affinity among the tested monosaccharides, it can be explored further for its therapeutic effects through *in silico* approaches.

**Table 3.** Binding affinity of monosaccharides of Spirulina polysaccharide to galectin-3.

| **Monosaccharide** | **Binding energy**<br>**(Kcal/mol)** | **Dissociation**<br>**constant(Ki)-mM** | **Interacting residues** |
| --- | --- | --- | --- |
| Galactose | -3.08 | 5.50 | Trp181A/H, Asn180A, Asn179A |
| Mannose | -2.75 | 9.59 | Trp181A/H, Asn180A, Gly182H |
| Rhamnose | -2.41 | 17.20 | Trp181B/F, Asn179B |
| Arabinose | -1.78 | 49.66 | Asn180B/F, Trp181B/F |
| Xylose | -1.63 | 64.14 | Trp181B/F, Asn179B, Asn180B |

**Figure 2.**
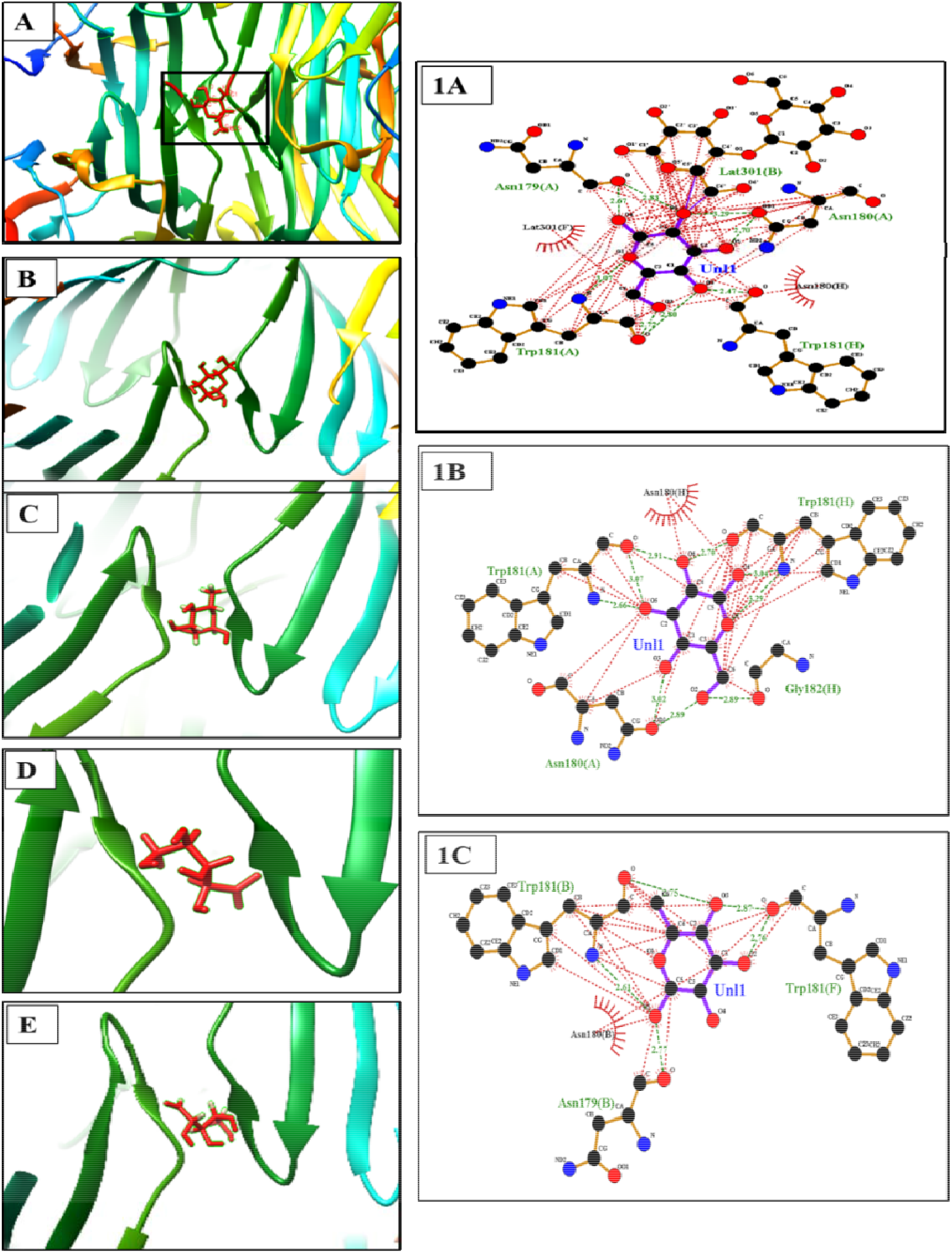
Interaction of Sp monosaccharides with Galectin-3. (A), (B), (C), (D), and (E) represent the binding pocket of Galactose, Mannose, Rhamnose, arabinose, and Xylose respectively with Galectin-3 protein. 1A, 1B, and 1C, are the hydrogen/hydrophobic interaction of A, B, and C with amino acid residues of galectin-3.

### 3.3 Cyto/DNA protection assays

The cellular protective ability of phytomolecules is particularly important as they regulate the cellular homeostasis and the downstream functions associated with it. we evaluated for the *in vitro* cytoprotective and DNA protective effects using oxidants like hydrogen peroxide, UV irradiations.

#### Buccal cell protection

The *in vitro* cytoprotective ability of Sp was tested on buccal cells against UV damage and the results were obtained as images, and observed for cellular damage including irregularities in cell shape. The fluorescent micrographs obtained after exposing to UV and treatments with gallic acid and Sp (Fig. 3), clearly showed that the Sp exhibited protective activity on buccal cells. Cells exposed to UV for different time intervals (10, 20, and 30 min) showed the appearance of clustering and cell disruption, while Sp treated samples almost regained their structural integrity revealing the cytoprotective effect of Sp in comparison to control and positive control like gallic acid.

**Figure 3.**
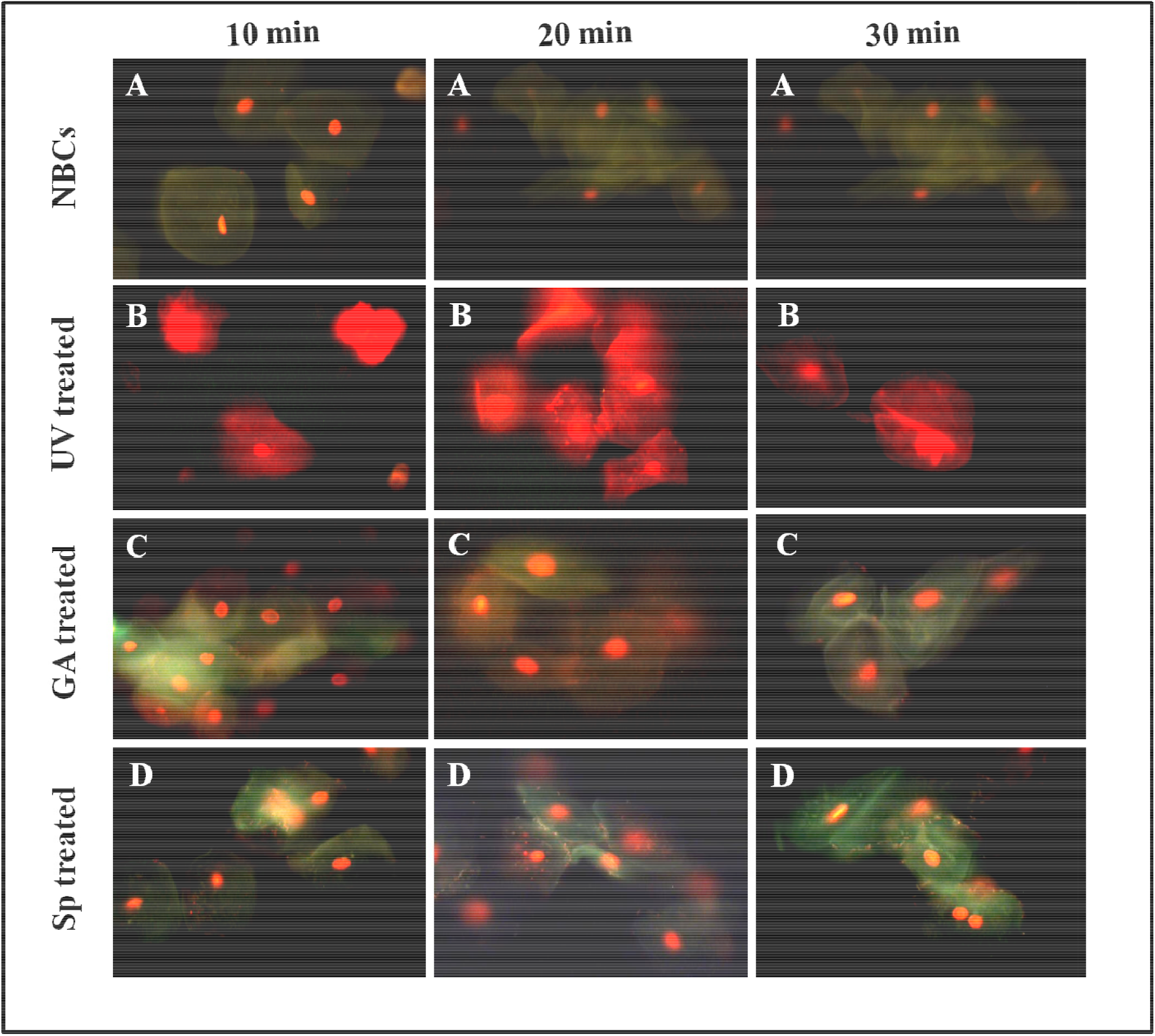
Buccal cells protection assay. Cell damage was induced on normal buccal cells (NBCs) by the exposure of cells to UV at 10, 20 and 30 min intervals, followed by the modulatory effect of the test compounds. (A) PBS control buccal cells, (B) UV treated buccal cells, and followed with the treatment of (C) gallic acid, (D) Spirulina polysaccharide. The intensity of red colour is proportional to cell death, and the green colour indicates the protective effects from UV damage.

#### In vitro RBCs (erythrocytes) protection assay

Since, RBCs are fragile and exposure to oxidants resulting in lysis, in this study we treated the cells with Sp and evaluated for the protective role. Briefly, the treatment with Sp prevented the RBCs from the damaging effects of oxidant and prevented the release of haemoglobin pigment into the medium. The absorbance at 410 nm is measured which indicates the extent of oxidative damage in RBCs. The treatment with Sp inhibited the haemolysis in a dose-dependent manner, and exhibited an IC_50_ value of 6.08 ± 0.05 μg/mL (Table 1), which is less when compared to standard gallic acid (9.02 ± 0.013 μg/mL) suggested a superior cellular protective ability by 32 %. Further, the scanning electron microscope (SEM) images upon treatment with Sp and exposure to hydrogen peroxide (H_2_O_2_) on RBCs structural morphology (cell rigidity and shape) were observed. The images showed clear morphological changes between the H_2_O_2_ exposure and treatments (Fig. 4). However, the Sp treatment prevented the change in shape upon H_2_O_2_ exposure similar to that of gallic acid treated cells confirms its cytoprotective effects.

**Figure 4.**
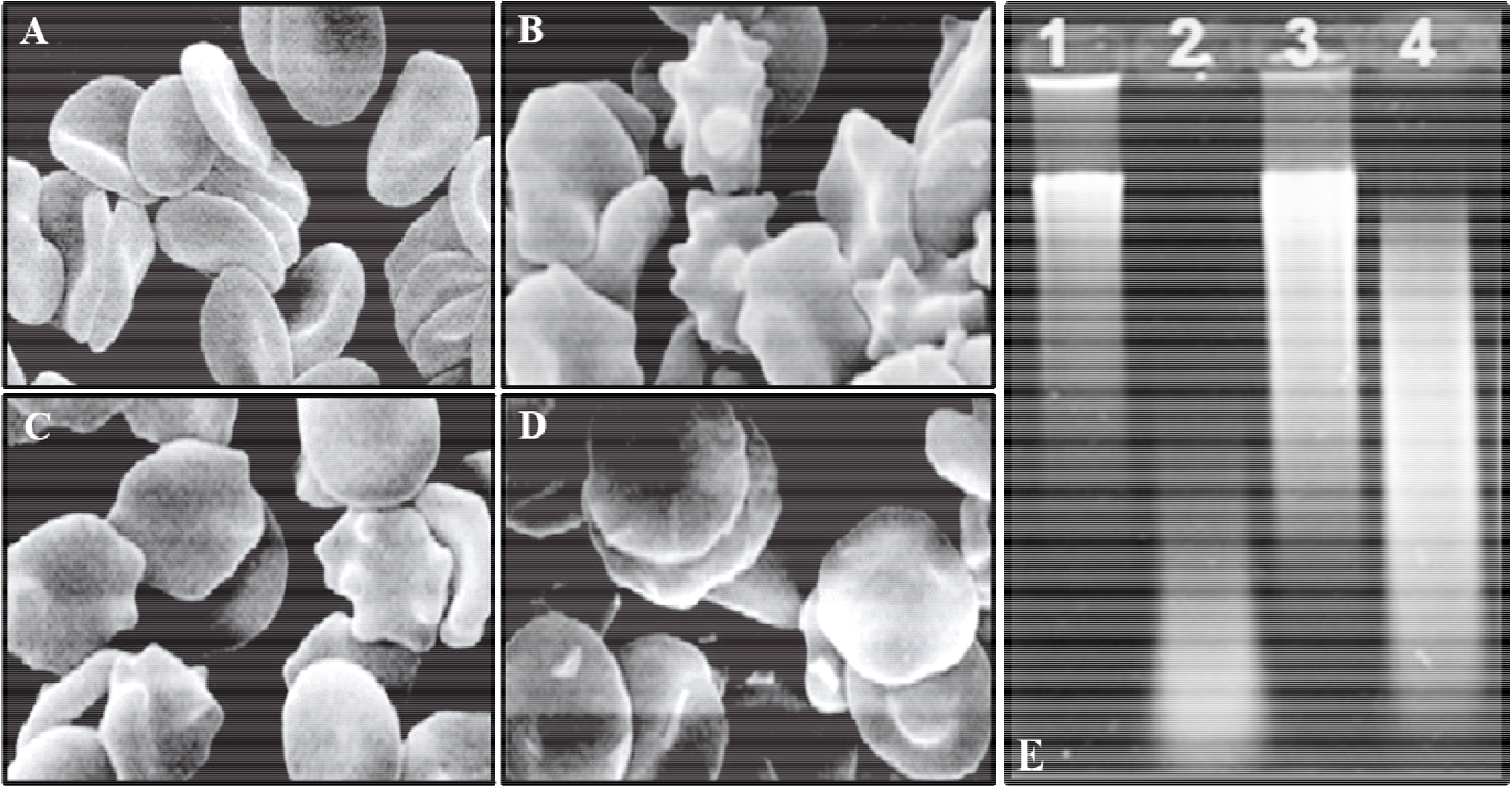
Cyto (RBCs) and DNA protective ability of Sp. (A) PBS control RBCs, (B) oxidant (H_2_O_2_) treated RBCs, (C) gallic acid treated RBCs, (D) Sp treated RBCs and (E) relative migration of DNA. Control DNA (1), oxidant treated DNA (2), followed with treatment of ascorbic acid (3), and Sp (4) on 1% Agarose gel. The relative migration between the lanes is compared, and analysed for the DNA protective effect of Sp. RBC: Red blood corpuscles; DNA: Deoxyribonucleic acid; PBS: Phosphate buffer saline; Sp: Spi ulina Polysaccharide.

#### DNA damage protection assay

The efficiency of Sp in preventing oxidative damage on calf thymus DNA was examined and evaluated for the migration pattern of treated and control cells (Fig. 4E). It was seen evidently that the native DNA (lane1) had lesser migration on the gel and incubation of DNA with H_2_O_2_ showed a significant decrease in the band intensity and had faster mobility suggesting degradation (lane 2) compared to control and treatments. However, pre-incubation of DNA with Sp (lane 4) prevented the DNA damage leading to the lower migration, in contrast to treatment with H_2_O_2_. Further, the DNA protective effects of Sp are comparable to gallic acid treated lane (lane 3), thus suggest the DNA protection ability of Sp. The observed DNA protective effect of Sp may be due to presence of phenolic compounds and associated antioxidant activity [Gargouri, et al., 2020; Liu, et al., 2011].

### 3.5 *In vitro* evaluation of Sp anticancer activity

#### Inhibitory effect of Sp on galectin-3 mediated RBCs agglutination and expression

Since, galectin-3 via its binding to β-galactosidases transforms normal cells to cancer cells, we evaluated the galectin-3 inhibitory activity of Sp in the *in vitro* system. We employed haemagglutination inhibition test as a measure for inhibition of metastasis as it has been accepted by several cancer laboratories (Sathisha et al., 2007; Gao, et al., 2013) including our laboratory. The MIC value of Sp was 6.25 μg/mL, which was 50 % higher activity than the standard galactose 9.37 μg/mL. Since, previous studies reported that the C-6 hydroxyl group of Galactose residues is responsible for higher affinity interaction with galactin-3 (Hirabayashi et al., 2002), the MIC data of Sp suggest that the observed inhibitory effect could be due blocking of this interaction. Similarly, the expression of galectin-3 and effect of Sp on its inhibition was studied on gastric cancer cells along with the standard commercial drug doxorubicin. The dose was fixed after the MTT assay, and AGS cells were treated with Sp (40 μg/mL) and the drug doxorubicin (40 μg/mL). The Sp treatment lowered the expression of galectin-3 significantly (p<0.05) by 38 % and doxorubicin by 53 % when compared to control (Fig. 5C). Thus, Sp showed potent galectin-3 inhibitory activity which is higher than the standard, thereby block or delay the process of metastasis in gastric cancer.

**Figure 5.**
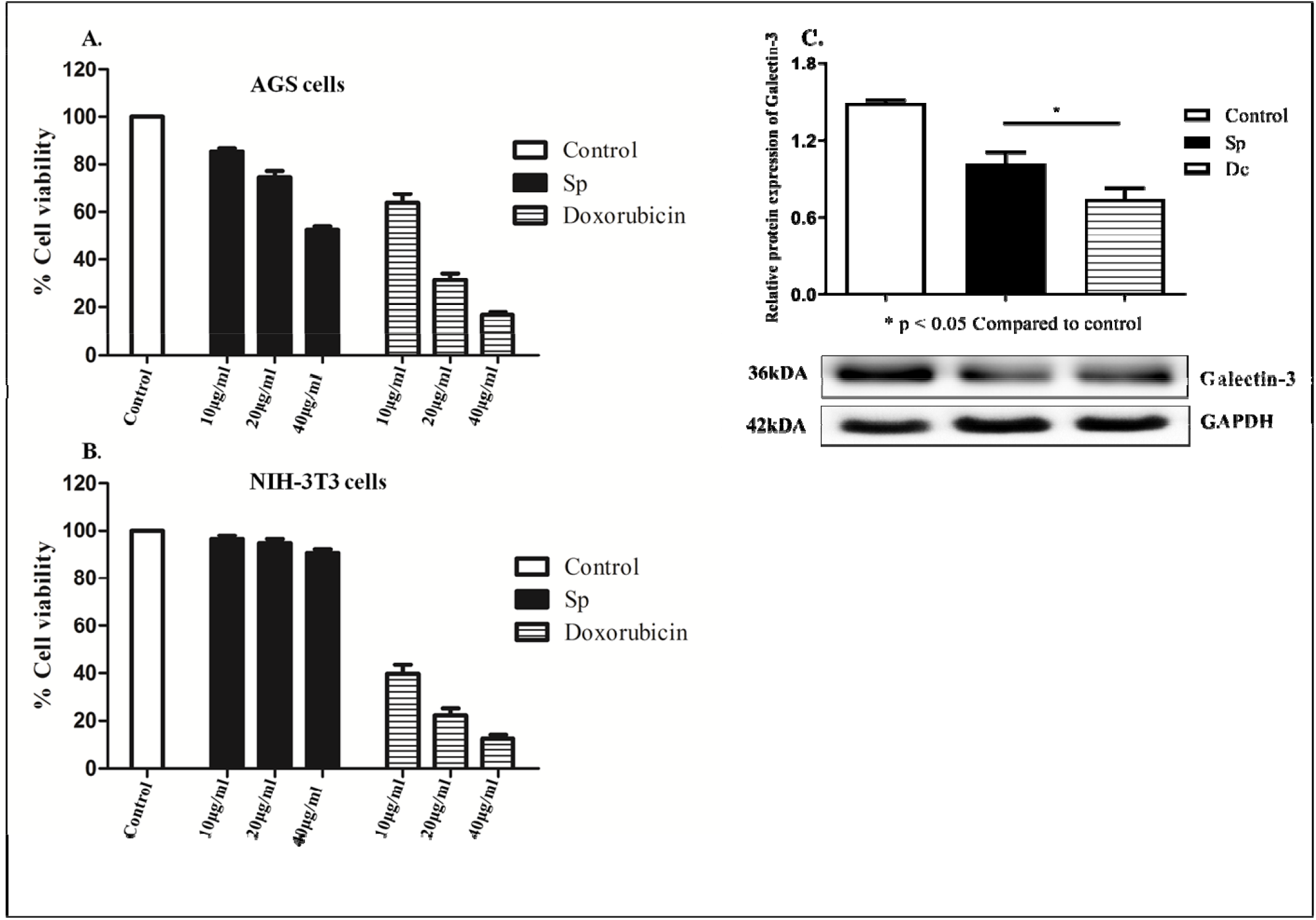
Antiproliferative effect of Sp on (A) AGS cells, and (B) NIH/3T3 cells, at different concentr tions (10, 20 & 40_μ_g/mL) in comparison to doxorubicin. (C) Relative protein expression of galactin-3 upon treatment with Sp and doxorubicin on AGS cells at similar dose (40_μ_g/mL). Values are mean ± SD of 3 samples and p value <0.05 is considered as statistically significant. AGS: Human Adenocarcinoma cells.

#### Antiproliferative assay

The gastric cancer cells were treated with Sp and doxorubicin at various concentrations and observed for the proliferation inhibition. Results depicted that Sp treatment at dose of 10, 20, and 40 μg/mL inhibited the proliferation of AGS cells by ∼15 to 48 %, and doxorubicin inhibited by ∼58 to 84 % at a similar concentration (Fig. 5A). Further treatments on NIH/3T3 normal cells revealed that the drug (doxorubicin) prevented the cells proliferation (80 %) by not just limiting to cancer cells, but by leading to the death of normal NIH/3T3 cells as well. The cytotoxic effects of doxorubicin observed in our study are in line with previous reports [Carvalho, et al., 2009; Damodar, et al., 2014] of clinical samples, whereas Sp treatment had a no major inhibition effect on the NIH/3T3 cells at the similar dose required for killing cancer cells. Thus, in spite of therapeutic effects of doxorubicin in killing AGS gastric cancer cells, its specificity remains as question that limits the applications. Meanwhile, with the advancement in the research on gastric cancer, now there is need for alternatives that precisely kills cancer cells with no or less effects on normal cells. In this approach, our results of Sp may be preferred over the drug due to its precise antiproliferative effects on AGS gastric cancer cells.

#### Apoptosis inducing ability of Sp

The AGS Cells were treated with Sp and doxorubicin, stained with Et-Br and acridine orange dye for the qualitative analysis of apoptosis. The cells were observed for the changes in the morphology of cytoplasm and chromatin condensation. The viable cells observed as green colour and dead cells nuclei appeared red colour. The Sp and doxorubicin treatment (40 μg/mL) significantly decreased the number of green colour stained cells. However, the drug showed more efficiency in causing cell death and inhibiting its proliferation than the Sp (Fig. 6C & 5A). The cause for the reduction in the proliferation of gastric cancer cells after treating with Sp and drug was further evaluated by DAPI staining and the data is represented in Fig. 6. Control cells had intact homologous nuclei and exhibited a little blue fluorescence, while the Sp and drug treated cells showed blue higher fluorescence of DAPI, thus indicating the loss of intact nuclei (Fig. 6). However, it was observed from fig. 6C, 6D, and 6E that Sp treated cells showed condensed nuclei along with apoptotic bodies similar to that of doxorubicin treatment.

**Figure 6.**
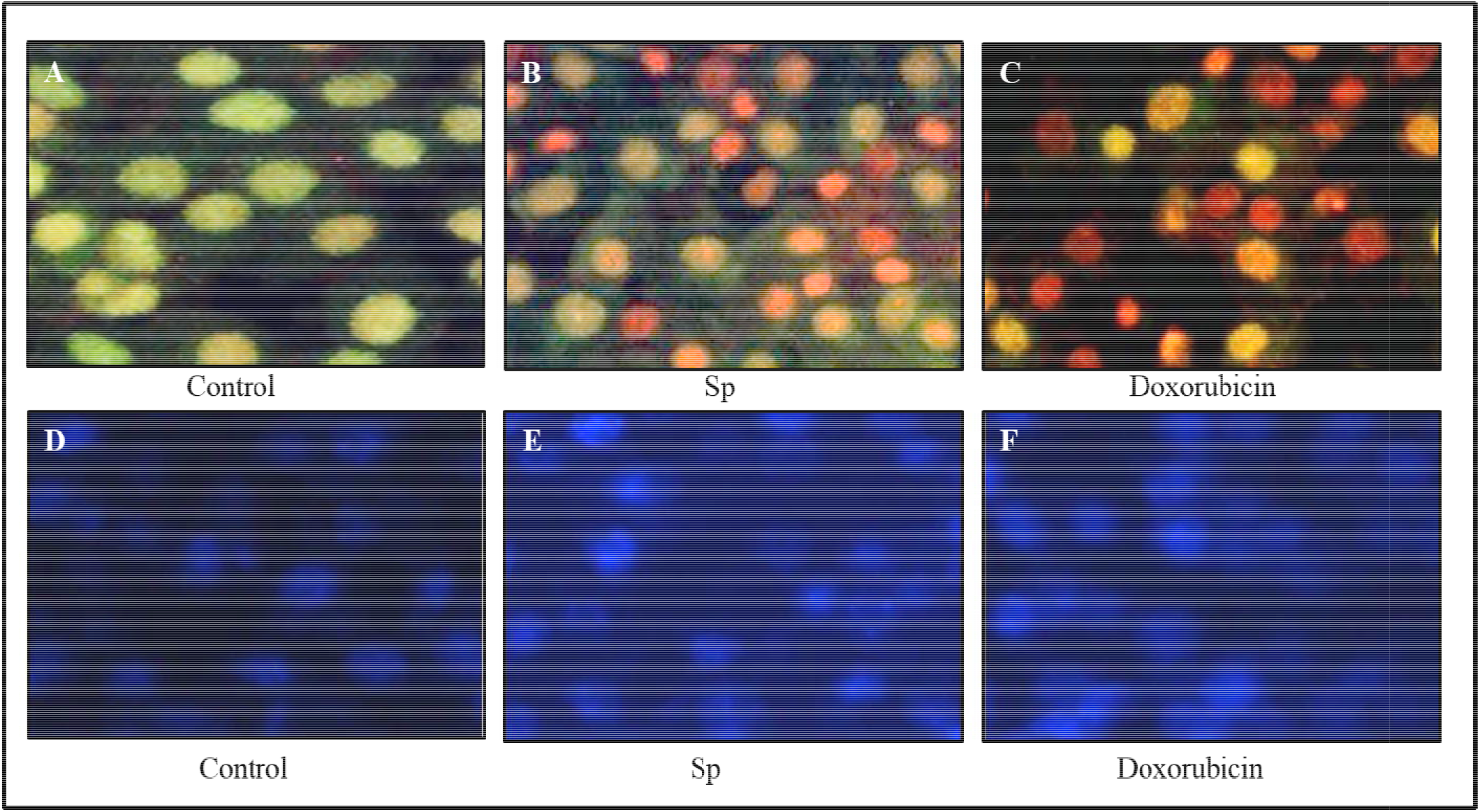
Effect of Sp on induction of apoptosis. Acridine orange and Et-Br stained images of (A) control, Sp, and (C) doxorubicin. DAPI stained images of (D), control, (E), Sp, and (F) doxorubicin treated AGS cells. Et-Br: Ethidium bromide; DAPI: 4, 6-diamidino-2-phenylindole; Sp: Spirulina polysaccharide.

#### In vitro tyrosinase inhibition of Sp

Since, the inhibition of tyrosinase a rate limiting enzyme in the synthesis of melanin have a clinical significance, we tested for the effect of Sp on the enzyme in the *in vitro* system. Surprisingly, Sp showed significant inhibition with IC_50_ value of 29.56 ± 3.85 μg/mL. Whereas, the synthetic inhibitor of tyrosinase like kojic acid had IC_50_ of 36.28 ± 4.12 μg/mL (Table 1). The previous studies reported the tyrosinase inhibitory effect of carotenoids like astaxanthin from green algae [Rao, et al., 2013], however there are no reports available for the effect of Sp on tyrosinase enzyme activity. Thus, this data strongly suggest the possibility of exploring the anticancer properties of Sp on skin cancer cells.

## 4. Discussion

Phytomolecules with antioxidant and anticancerous properties offer a safe strategy to prevent the onset of various types of cancer. Many epidemiological studies found that the risk of gastric cancer was inversely associated with the intake of natural products with therapeutic potentials [Fang, et al., 2015]. The spread of cancer could be directly attributed to the galectin-3 marker, and the carbohydrates specifically polysaccharides are under constant scrutiny [Yang, et al., 2008]. Interestingly, the action of galectin-3 mediated interactions is vital on the carbohydrate-binding domain of galectin-3 between the normal and cancer cells [Khan, et al., 2019]. Therefore, the dietary oligosaccharides or polysaccharides garnering the attention and the supremacy lays in the binding ability of specific sugars to block the binding of galectin-3 to the extracellular matrix of the adjacent cells which otherwise cause metastasis. [Jayaram, et al., 2015; Ahmed, et al., 2015]. Various investigators optimised the extraction conditions and partially characterized the sulphated Sp [Rajasekar, et al., 2019] and explored for the potential antioxidant properties in *in vitro* models [Kurd, et al., 2015]. It has been shown that sulphated spirulina polysaccharides exhibited the greater antibacterial activity and improved the reproductive performance in zebra fish [Rajasekar, et al., 2019]. However, the anticancerous effect of Sp on the cancer cells is rarely explored. Therefore, we specifically targeted the gastric cancer cells to assess the anti-cancerous activity of Sp as gastric cancer is the the third most prevalent cause of cancer-related deaths in the world and the dietary polysaccharides are known to exhibit the anti-metastatic properties. This study demonstrated the *in vitro* anti-cancerous properties of the Sp by assessing its ability to intervene the tumour specific marker galectin-3 in gastric cancer cells, along with the elucidation of possible structure to function relationship evidence.

Spirulina is considered as health supplement throughout the world and the beneficial effects of spirulina whole biomass, components like phycocyanin, carotenoids are well reported. Although antiviral properties of sulphated polysaccharides are mentioned in the literature, their role in the cancer prevention / treatment is not documented. In this direction the present results may pave the way for further studies of spirulina polysaccharides in cancer prevention without sideouts unlike the cancer drugs in use. The evidence from this study indicated that Sp possesses average high molecular weight of 1457 kDA, and the presence of galactoarabinorhamnoglycan backbone. Also, moderate correlation was observed between arabinose and galactose (p>0.05) [data not shown], and contrastingly, no association was seen for the other sugars of Sp. It was observed that a known anticancerous drug doxorubicin inhibited the normal (NIH/3T3) cells growth in addition to the inhibition of AGS cell growth (Fig. 5), clearly indicating the toxic effect of doxorubicin. Therefore, the Sp which specifically inhibited the proliferation of gastric cancer cells without affecting the normal cells may be preferred for better therapeutic approaches. Further, the Sp treatment on AGS cells showed the induction of apoptosis as evidenced by stained images with Et-Br/acridine orange and DAPI suggests the possible underlying mechanism. Similarly, the Sp significantly (p<0.05) lowered the galectin-3 mediated agglutination formation and expression in AGS cells confirm the anti-metastatic activity is by modulation of galectin-3. The i*n silico* interaction study supports the findings that galactose is the major sugar interacting with galectin-3 protein and predicts that Sp may exhibit anticancer property by inhibiting galectin-3. In this study we observed the protective effects of Sp on RBCs, buccal cells, and DNA, against oxidant damage strongly suggest the cytoprotective effects in addition to its antioxidant properties. Since, the use of blue green algae like spirulina is escalating in modern times and because of its nutritional composition and biomolecules that it possesses, the evidence on the bioactive properties will diversify its currently known applications.

In conclusion, the current study addresses the first line report of the galectin-3 inhibitory potential of Sp and its effect on AGS gastric cancer cells as opposed to commercially available synthetic drug doxorubicin. The structural characterization data presented gives the possible reasons for the observed antioxidant and anti-metastatic property of Sp. The cytoprotective effects of Sp open the possibility of exploring its use in oxidative stress and inflammation induced diseases as well. Although, the study addressed the effect of Sp on galectin-3 modulatory potential and the structural evidence for the possible effects, the futuristic research can be taken as follows. (1) derivatising to oligosaccharides as there are few reports that suggests oligos have a better effect than polysaccharides, (2) evaluation of the effect of Sp in other cancer cell types, (3) attempts to see the synergistic effects with currently available drugs, and (4) elucidating the mechanism of action and bioavailability in *in vivo* animal models will diversify the currently known medicinal, nutraceutical, nutritional, and food application of the blue green algae.

## Abbreviations

Sp: spirulina polysaccharide
Vit.c: Vitamin C
RBC: Red blood corpuscles
MTT: 3-[4,5-dimethyl-2-thiazolyl]-2,5-diphenyl 2-H-tetrazolium bromide
DNA: deoxyribonucleic acid
UV: ultraviolet
IC_50_: Inhibition concentration 50
AGS: Adenocarcinoma gastric cell line
CAE: cellulose acetate electrophoresis
NMR: Nuclear magnetic resonance
FTIR: Fourier transfer ionization infrared radiation
GAE: Gallic acid equivalent
DAPI: 4′,6-diamidino-2-phenylindole
Et-Br: Ethidium Bromide
TEMED: Tetramethylethylenediamine

---

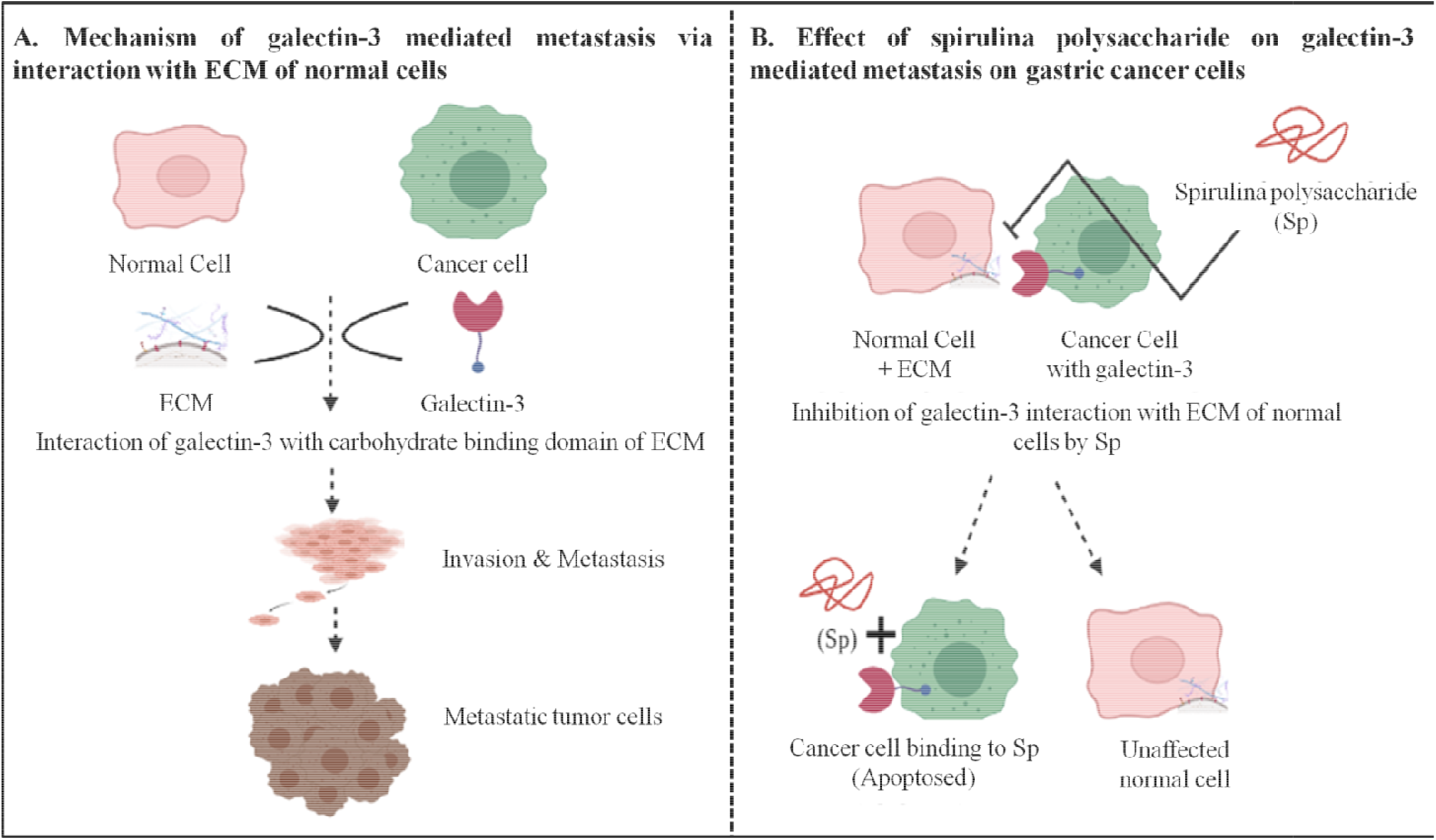
Graphical representation of mechanism of galectin-3 mediated metastasis and intervention by spirulina polysaccharide. The interaction between galectin-3, a molecule on the surface of cancer cells with extracellular matrix of the surrounding normal cells is believed to be the reason for the metastasis. The galactose residue of the Sp interaction with galectin-3 is believed to possess galectin inhibitory property are the same has been tested. Sp intervene this interaction and thus preventing cancer metastasis. ECM Extracellular Matrix; Sp: Spirulina Polysaccharide.

## Highlights

➢ Spirulina polysaccharide was isolated by hot water reflux method, purified and partially characterized.
➢ The bioactive properties such as cyto/DNA protection, Anti-metastasis, and Anti-cancer properties were evaluated.
➢ The study specifically evaluated the galectin-3 modulatory potential of spirulina polysaccharide (Sp), a universal biomarker for all types of metastatic cancers.
➢ Sp could be used as a potential galectin-3 inhibitor in the functional foods.

